# A self-immolative linker for heparanase activatable probes

**DOI:** 10.1101/2021.06.15.448502

**Authors:** Kelton A. Schleyer, Jun Liu, Zhishen Wang, Lina Cui

**Affiliations:** Department of Medicinal Chemistry, College of Pharmacy, UF Health Science Center, UF Health Cancer Center, University of Florida, Gainesville, FL 32610, USA

## Abstract

Substrate-based probes utilize known substrate specificity parameters to create a probe that can be activated by a target enzyme. In developing probes for heparanase, an endo-ß-glucuronidase, we previously reported that small, inactive substrate-based probes could be electronically tuned by incorporating electron-withdrawing atoms on the aromatic aglycone fluorophore, *ortho*- to the cleaved glycosidic bond. However, the installation of electron-withdrawing groups directly onto established fluorophores or other reporters complicates the synthesis of new heparanase probes. In this work we report a new design strategy to expand the toolkit of heparanase imaging probes, in which the installation of an electronically tuned benzyl alcohol linker restored the activity of a previously inactive heparanase probe using 4-methylumbelliferone as the fluorescent reporter, suggesting such a linker can provide a scaffold for facile development of activatable heparanase probes bearing a variety of imaging moieties.

## Maintext

Heparanase (HPSE) is an endo-β-glucuronidase that cleaves glycosidic bonds of heparan sulfate (HS) of heparan sulfate proteoglycans (HSPG) localized in the extracellular matrix and basement membrane.^1^ As HS plays critical roles in maintenance of integrity of the ECM and modulation of the activity of diverse, cytokines, chemokines and growth factors, the degradation of HS mediated by heparanase affected a variety of biological processes.^2–4^ Under normal physiological conditions, high-level heparanase can be detected in the placenta and blood-borne cells such as platelets, neutrophils, mast cells, and lymphocytes.^3^ Recently, the data revealed that heparanase is overexpressed in tumor tissues^1, 5–7^ and the enzymatic activity is correlated with tumor metastasis, angiogenesis and post-surgical survival.^8^ Besides, the role of this enzyme is also associated with inflammatory disorders^3, 9^ and autoimmune diabetes.^10–11^ Thus, it is unsurprising that heparanase has been regarded as a promising pharmacological target for treatment of multiple diseases.^3–4, 12–14^

Recently, we reported the first structurally defined ultrasensitive fluorogenic probe **HADP (1)** for detecting heparanase activity with high selectivity and sensitivity, which we used in a high-throughput screen for novel heparanase inhibitors.^15^ In our initial probe design, use of the fluorogenic reporter 4-methylumbelliferone (4-MU) did not facilitate turnover of the probe by HPSE, consistent with the exclusively *endo*-glycosidic selectivity of HPSE enzymatic activity. However, by incorporating two electron-withdrawing fluorine atoms on the methylumbelliferone reporter, *ortho* to the phenolic oxygen, our **HADP** probe successfully elicited *exo*-glycosidic activity from HPSE.^15^ Following this study, we developed a second fluorogenic probe for imaging heparanase activity in living cells (J.L., Z.W., and L.C., unpublished results), which also required two *ortho*-position fluorine atoms installed on the reporter, to elicit HPSE activity. Encouraged by these advances, we attempted to develop a near-infrared (NIR) probe for monitoring heparanase *in vivo* by extending the conjugation system of fluorophore, but the synthesis failed due to incompatibility of the NIR fluorophore with the chemical manipulation of the disaccharide recognition unit. Thus, we considered adding a self-immolative linker between the recognition unit and fluorescent reporter unit (**Fig. 1**). Typically, a self-immolative linker is based on a 4-hydroxybenzyl alcohol moiety to bridge the recognition unit and fluorophore.^16–17^ Once the recognition unit reacts with the corresponding analyte, the phenol of the middle linker is uncaged, which triggers simultaneous elimination of a molecule of para quinone methide to release the latent reporter unit. This strategy has been extensively employed in the design of prodrugs, drug delivery, sensors, and molecular amplifiers. This self-immolative linker can efficiently reduce steric limitations and increase stability. Our previous work showed that *o,o*-difluorination of the fluorophore aglycone elicited heparanase activity while a non-fluorinated analog did not. This was due to the resultant weakening of the glycosidic bond and lowering of the transition state energy for enzymatic turnover by heparanase.^15^ Based on these results, we deigned two molecules bearing a 4-hydroxybenzyl alcohol linker substituted with either two or four fluorine atoms to facilitate enzymatic cleavage by heparanase. A simple coumarin as fluorescent reporter to examine whether the linker can be disassembled upon incubation with heparanase. Our results show that a tetrafluorobenzaldehyde (**4-F**) linker facilitates heparanase turnover, possibly providing a universal scaffold for small molecule heparanase probes.

**Fig. 1.**
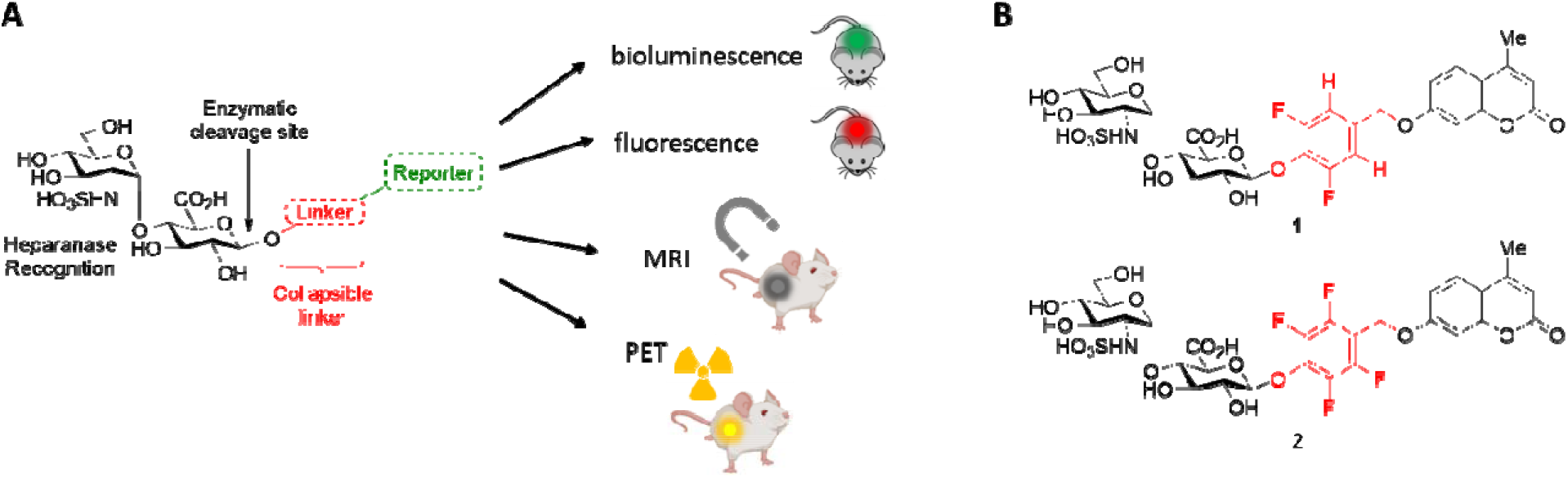
**(A)** Design of HPSE probes bearing self-immolative linkers. This will serve as a scaffold for probes of many imaging modalities. **(B)** Probes **1** and **2** designed in this work, using differently tuned linkers.

We embarked on the synthesis for compounds **1** and **2** (**Scheme 1**) from disaccharide bromide **3** with 4-hydroxybenzaldehyde derivatives bearing 2 or 4 fluorine atoms by Koenigs-Knorr glycosylation to afford compounds **6** and **7**, respectively. The aldehyde group was subsequently reduced to the primary alcohol, followed by an Appel reaction to convert the alcohols to benzyl bromides **10** and **11**. The alkyl bromides were substituted with 4-methylumbelliferone in the presence of potassium carbonate to incorporate the fluorescent reporter, establishing the skeleton of the desired molecules (compounds **12** and **13**). Subsequent deprotection steps included Zemplén *O*-deacylation, catalytic hydrogenation of the azido group to the respective amine, and saponification of the glucuronic acid methyl ester. Finally, glucosamine *N*-sulfation afforded compounds **1** and **2**.

**Scheme 1.**
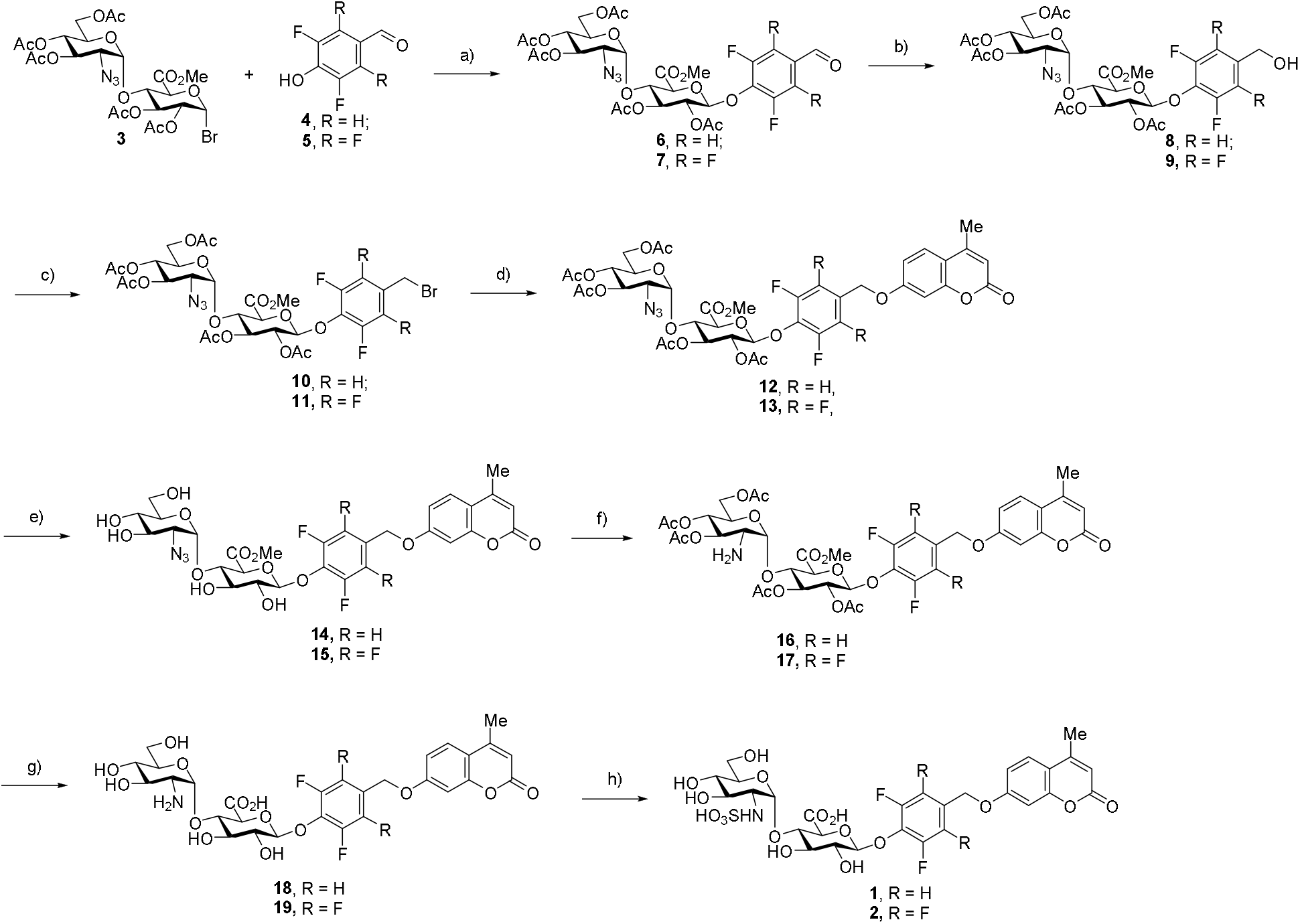
Synthetic scheme for probes **1** and **2**. Reagents and conditions: a) Ag_2_O, MeCN (dry), r.t., OVN, **77%** (**6**) or **54%** (**7**); b) NaBH_4_, DCM/MeOH (1/5), r.t., 20 min, **49%** (**8**) or **64%** (**9**); c) PPh_3_, CBr_4_, DCM (dry), r.t., 1 h, **46%** (**10**) or **72%** (**11**); d) 4-MU, K_2_CO_3_, DMF, r.t., 16 h, **81%** (**12**) or **47%** (**13**); e) NaOMe, MeOH, r.t., 2 h, **quant.** (HPLC); f) Pd/C, H_2_, EtOAc, rt, 18 h, **75%** (HPLC); g) NaOH (pH 12), H_2_O, rt, 4 h, **quant.** (HPLC); h) Py·SO_3_, NaOH (pH 10), H_2_O, rt, 6 h, 50% (HPLC).

To confirm that the linkers work, we investigated the enzymatic response of compounds 1 and 2 to heparanase in NaOAc buffer (pH 5.0) by measuring fluorescence turn-on from the (putatively) release methylumbelliferone reporter. For compound 1 with two fluorine atoms, negligible fluorescence was observed after incubation for 24 hours, indicating this linker could not be cleaved by heparanase.

Furthermore, the reaction solution of this compound with heparanase was injected to HPLC. it shows that predominant peak remained, and the retention time was consistent with that of the compound 1. In contrast, compound 2 displayed a dramatic fluorescence enhancement after incubation for 3 hours in NaOAc buffer, at pH 5.0 (**Fig. 2A**). When the assay was terminated by increasing the pH to 10, the turn-on ratio can be further boosted even with a diluted mixture (**Fig. 2B**). This cleavage is further confirmed by HPLC trace, giving a new peak in alignment with the corresponding free 4-methylumbelliferone in a complete conversion (**Fig. 3**).

**Fig. 2.**
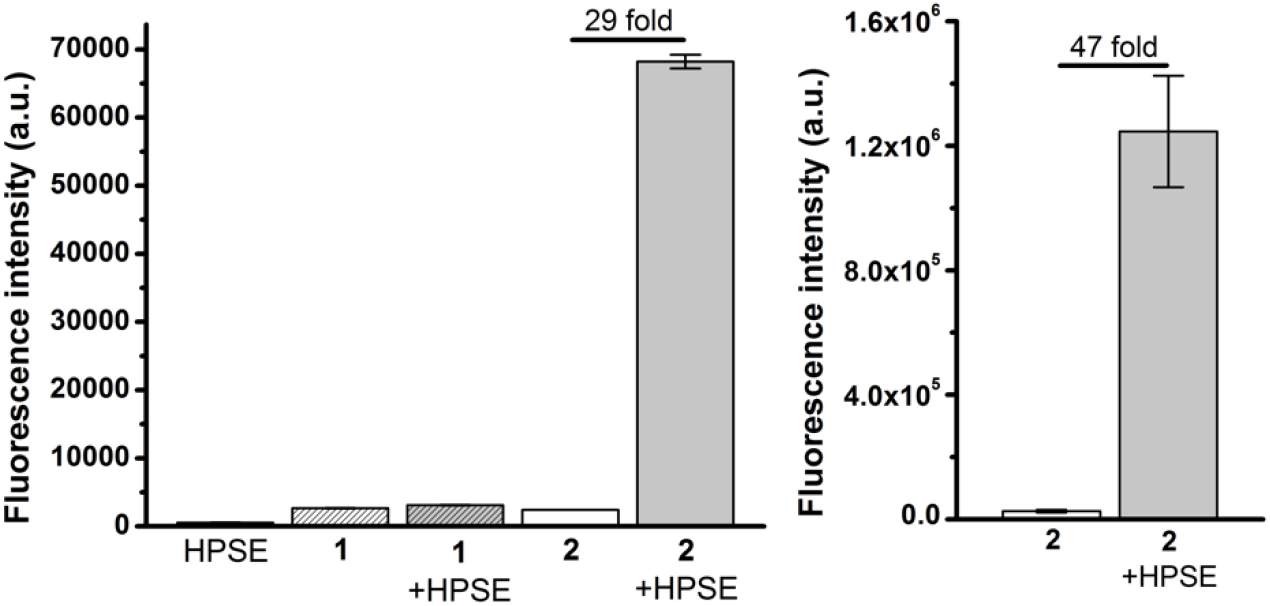
Fluorescence emission of **1** and **2** upon incubation with HPSE after 24 h. (**B**) Fluorescence of **2** after 24 h incubation, adjusted to pH 10.

**Fig. 3.**
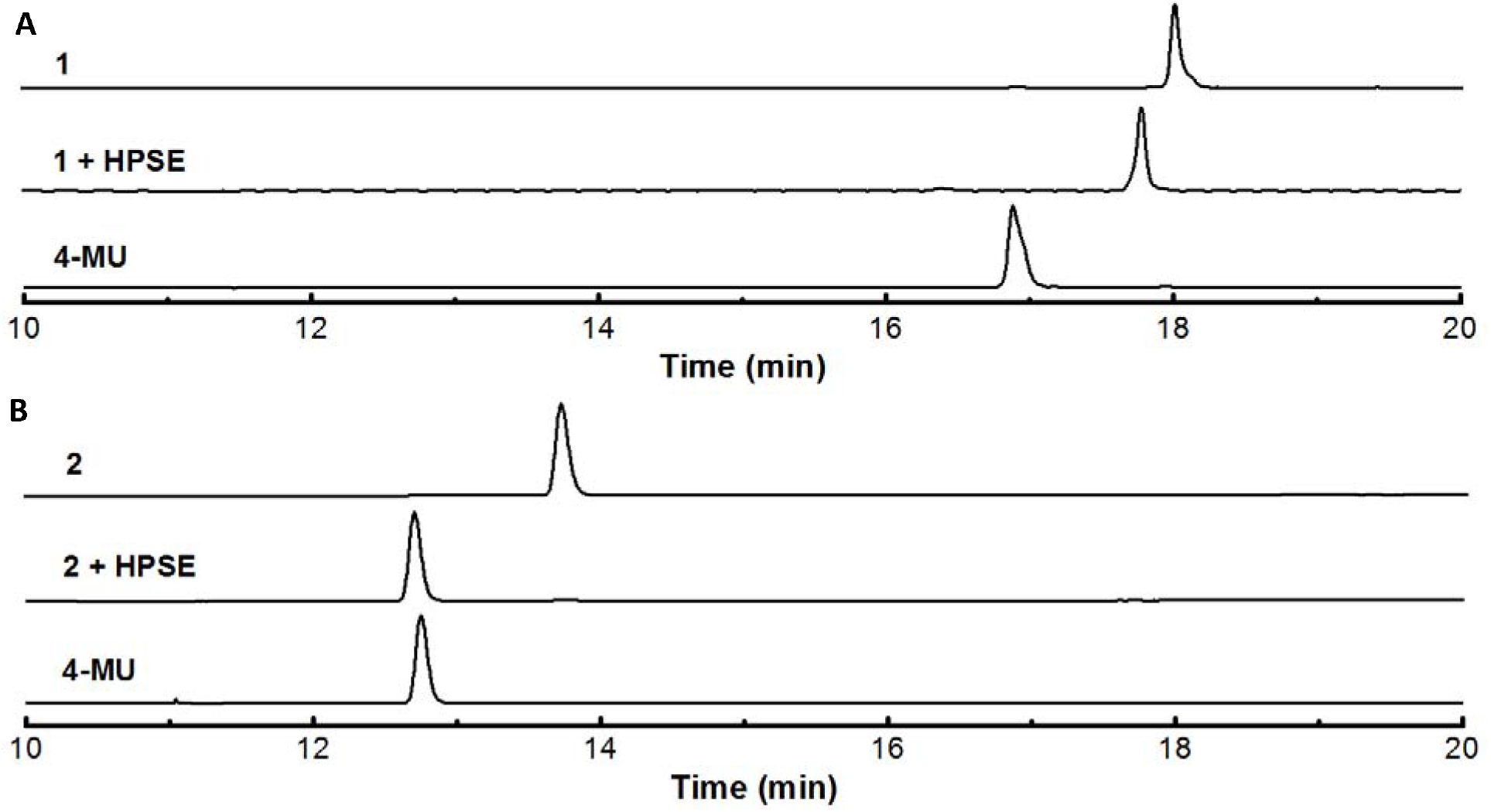
HPLC trace of (**A**) **2-F** or (**B**) **4-F** with or without HPSE after 24 h. Included is the free fluorophore, **4-MU**.

The successful response of **2** to HPSE suggests that the electronically tuned 4-F linker can facilitate turnover of probes using non-activated reporters, such as 7-MU. The 4-F scaffold thus opens up potential for the use of traditional reports in detecting HPSE activity, including commercially available fluorophores, chelates for magnetic or radioactive contrast, or smart imaging scaffolds, such as bioluminescence and *in situ* self-assembly moieties. The inclusion of 4-F avoids the requirement of functionalizing each of these reporters with EWGs such as the difluorination of our previously reported probe **HADP**, likely providing convenient and facile synthetic access to a library of novel HPSE imaging probes. We are currently synthesizing novel probes using the 4-F scaffold to expand this theory.

## Acknowledgements

This work is supported by research grants to Prof. L. Cui from the National Institute of General Medical Sciences of National Institutes of Health (Maximizing Investigators’ Research Award for Early Stage Investigators, R35GM124963), the Department of Defense (Congressionally Directed Medical Research Programs Career Development Award, W81XWH-17-1-0529), and to K.A. Schleyer from the University of Florida Health Cancer Center (UFHCC Predoctoral Award). We are grateful for the support from the University of Florida (UF) and the UF Health Cancer Center. NMR spectra were collected from the Department of Medicinal Chemistry, College of Pharmacy, University of Florida (UF). Mass spectrometry services were provided in part by the Mass Spectrometry Research and Education Center, Department of Chemistry, UF (supported by NIH S10 OD021758-01A1).

## Author contributions

JL and LC designed the project. KAS, JL, and ZW synthesized the compounds. JL and KAS performed the fluorescence assays and HPLC characterization. KAS and LC wrote the paper. All authors reviewed the manuscript.

## Conflict of Interest

The University of Florida has filed a patent on the work.

## Methods

Experimental methods and chemical characterization are provided in the Supplementary Information.

