## Supporting information for "A self-immolative linker for heparanase activatable probes"

### Chemical Synthesis

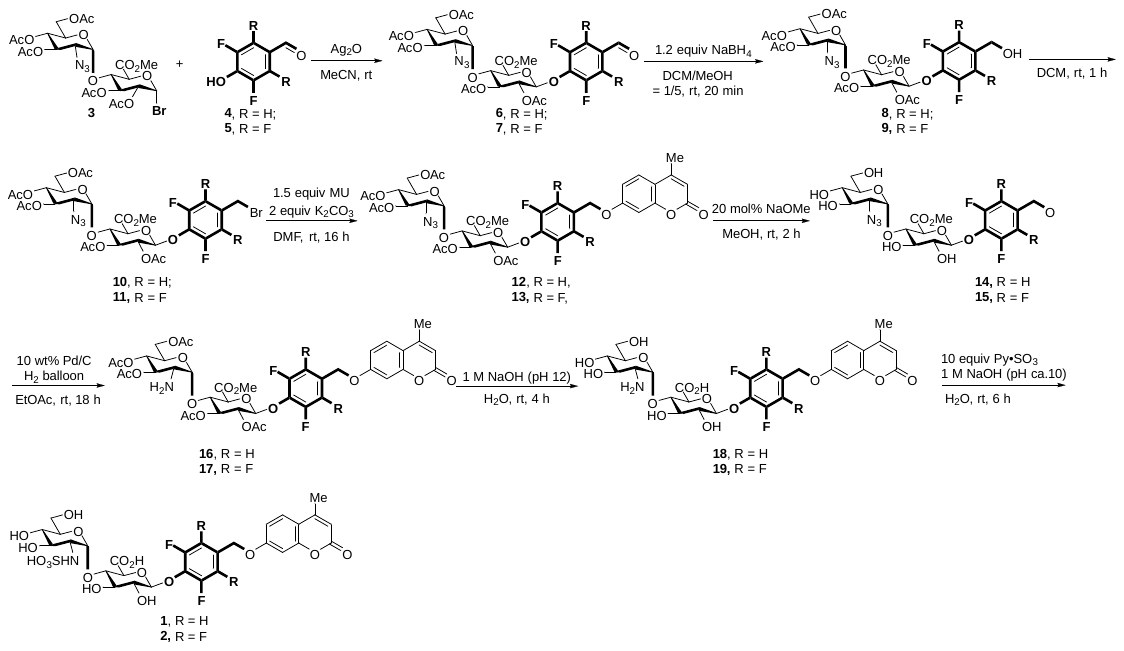
Scheme S1. Synthetic scheme for compounds used in this work.

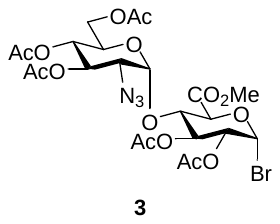

**Methyl (3’,4’,6’-Tri-O-acetyl-2’-azido-2’-deoxy-α-D-glucopyranosyl)-(1→4)-(1-bromo-1- deoxyl-2,3-di-O-acetyl-β-D-glucopyranosyluronate (3):** Compound **3** was synthesized according to our previously reported work.^1^

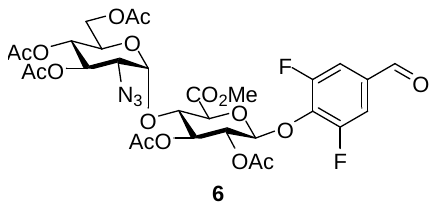

**Methyl (3’,4’,6’-Tri-O-acetyl-2’-azido-2’-deoxy-α-D-glucopyranosyl)-(1→4)-(1-(3,5-difluoro-4-hydroxybenzaldehyde)-1- deoxyl-2,3-di-O-acetyl-β-D-glucopyranosyluronate (6):** To a solution of **3** (125 mg, 0.187 mmol) in freshly distilled acetonitrile (2 mL) was added 3,5-difluoro-4-hydroxybenzaldehyde (35 mg, 0.224 mmol, 1.2 equiv.) and Ag_2_O (87 mg, 0.374 mmol, 2.0 equiv.). The reaction was covered in tin foil and stirred under argon at room temperature overnight. Conversion of product was monitored by TLC (R*_f_* = 0.34, hexanes/ethyl acetate 1:1). The reaction was filtered over Celite and concentrated *in vacuo*, then purified *via* silica gel chromatography using hexanes/ethyl acetate 2:1 then hexanes/ethyl acetate 1:1. (**77%**). ^1^H NMR (600 MHz, Chloroform-*d*) δ 9.88 (t, *J* = 1.6 Hz, 1H), 7.51 (d, *J* = 7.3 Hz, 2H), 5.44 – 5.37 (m, 2H), 5.33 (dd, *J* = 10.7, 9.3 Hz, 1H), 5.28 – 5.22 (m, 2H), 5.03 (dd, *J* = 10.3, 9.3 Hz, 1H), 4.41 (t, *J* = 8.8 Hz, 1H), 4.30 – 4.22 (m, 2H), 4.10 (dd, *J* = 12.5, 2.2 Hz, 1H), 3.87 (ddd, *J* = 10.3, 3.8, 2.3 Hz, 1H), 3.78 (s, 3H), 3.47 (dd, *J* = 10.7, 3.7 Hz, 1H), 2.11 (d, *J* = 5.1 Hz, 6H), 2.08 (d, *J* = 5.5 Hz, 6H), 2.03 (s, 3H). ^13^C NMR (151 MHz, CDCl_3_) δ 188.65, 170.35, 169.62, 169.39, 169.38, 169.30, 167.17, 156.22, 156.20, 154.54, 154.51, 137.26, 132.35, 113.28, 113.25, 113.16, 113.12, 100.75, 98.56, 75.26, 74.02, 72.85, 71.10, 69.86, 68.39, 67.88, 61.00, 60.67, 52.77, 20.43, 20.42, 20.37, 20.31, 20.29. HRMS (ESI) for C_30_H_33_F_2_N_3_O_17_ [M+Na]^+^ : Calc’d: 768.1670, Found: 768.1690.

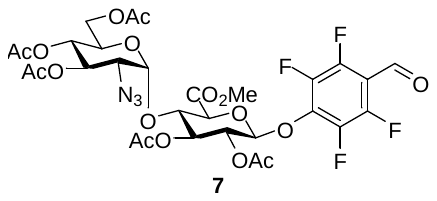

**Methyl (3’,4’,6’-Tri-O-acetyl-2’-azido-2’-deoxy-α-D-glucopyranosyl)-(1→4)-(1-(2,3,5,6-tetrafluoro-4-hydroxybenzaldehyde)-1- deoxyl-2,3-di-O-acetyl-β-D-glucopyranosyluronate** **(7):** To a solution of **3** (125 mg, 0.187 mmol) in freshly distilled acetonitrile (2 mL) was added 2,3,5,6-tetrafluoro-4-hydroxybenzaldehyde (44 mg, 0.224 mmol, 1.2 equiv.) and Ag_2_O (87 mg, 0.374 mmol, 2.0 equiv.). The reaction was covered in tin foil and stirred under argon at room temperature overnight. Conversion of product was monitored by TLC (R*_f_* = 0.57, hexanes/ethyl acetate 1:1). The reaction was filtered over Celite and concentrated *in vacuo*, then purified *via* silica gel chromatography using hexanes/ethyl acetate 2:1 then hexanes/ethyl acetate 1:1. (**54%**).^1^H NMR (600 MHz, Chloroform-*d*) δ 10.18 (s, 1H), 5.49 (d, *J* = 6.2 Hz, 1H), 5.32 – 5.23 (m, 2H), 5.21 – 5.13 (m, 2H), 4.95 (dd, *J* = 10.3, 9.3 Hz, 1H), 4.37 (t, *J* = 8.2 Hz, 1H), 4.23 (d, *J* = 8.4 Hz, 1H), 4.17 (dd, *J* = 12.5, 4.0 Hz, 1H), 4.08 – 4.00 (m, 1H), 3.82 (ddd, *J* = 10.3, 4.0, 2.2 Hz, 1H), 3.71 (s, 3H), 3.37 (dd, *J* = 10.8, 3.7 Hz, 1H), 2.06 (s, 3H), 2.04 (s, 3H), 2.01 (d, *J* = 2.1 Hz, 6H), 1.97 (s, 3H). ^13^C NMR (151 MHz, CDCl_3_) δ 181.97, 171.12, 170.55, 169.83, 169.57, 169.46, 169.41, 167.20, 148.27, 146.58, 146.54, 141.36, 141.27, 139.69, 139.60, 139.17, 139.09, 110.79, 110.73, 110.66, 100.30, 100.28, 100.26, 98.83, 75.05, 74.08, 72.19, 70.62, 70.03, 68.67, 68.04, 61.21, 60.84, 60.36, 53.04, 21.00, 20.59, 20.52, 20.50, 14.17. HRMS (ESI) for C_30_H_31_F_4_N_3_O_17_ [M+Na]^+^ : Calc’d: 804.1487, Found: 804.1499.

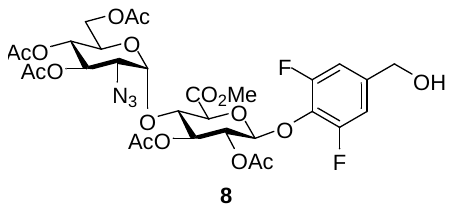

**Methyl (3’,4’,6’-Tri-O-acetyl-2’-azido-2’-deoxy-α-D-glucopyranosyl)-(1→4)-(1-(2,6-difluoro-1-hydroxy-4-methylhydroxybenzyl)-1- deoxyl-2,3-di-O-acetyl-β-D-glucopyranosyluronate (8):** To a solution of **6** (107 mg, 0.144 mmol) in freshly distilled DCM (2 mL) was added a solution of sodium borohydride (6.5 mg, 0.173 mmol, 1.2 equiv.) in dry MeOH (200 µL) and the reaction was stirred at room temperature. Conversion of product was monitored by TLC (R*_f_* = 0.43, hexanes/ethyl acetate 1:1). Upon completion, the reaction was quenched by addition of water and extracted into ethyl acetate. The organic layer was washed with brine and dried of Na_2_SO_4_, then purified by FCC using isocratic hexanes/ethyl acetate 1.5/1. White solid product acquired, 53 mg (**49%**). ^1^H NMR (600 MHz, Chloroform-*d*) δ 6.92 (d, *J* = 8.4 Hz, 2H), 5.29 (t, *J* = 9.0 Hz, 1H), 5.21 (dd, *J* = 9.0, 7.3 Hz, 1H), 5.05 (ddd, *J* = 19.1, 15.5, 7.2 Hz, 4H), 4.62 (d, *J* = 5.1 Hz, 2H), 4.57 (d, *J* = 10.2 Hz, 1H), 4.34 (t, *J* = 9.0 Hz, 1H), 4.18 (dd, *J* = 12.5, 3.6 Hz, 1H), 4.08 – 4.04 (m, 1H), 4.00 (d, *J* = 9.3 Hz, 1H), 3.96 (dt, *J* = 10.3, 5.2 Hz, 1H), 3.76 (s, 3H), 3.72 (dt, *J* = 9.9, 3.0 Hz, 1H), 2.25 – 2.19 (m, 1H), 2.10 – 2.04 (m, 9H), 1.98 (d, *J* = 2.2 Hz, 6H), 1.41 (s, 8H). ^13^C NMR (151 MHz, CDCl_3_) δ 171.34, 170.96, 170.92, 169.92, 169.67, 169.34, 167.38, 156.53, 156.50, 155.21, 154.86, 154.83, 139.14, 131.52, 110.37, 110.33, 110.24, 110.21, 102.10, 99.08, 80.39, 75.06, 74.81, 73.71, 71.58, 71.08, 68.98, 67.77, 63.73, 61.35, 60.53, 53.10, 53.08, 28.28, 21.16, 20.81, 20.73, 20.68, 20.66, 14.30. HRMS (ESI) for C_30_H_35_F_2_N_3_O_17_ [M+Na]^+^ : Calc’d: 770.1832, Found: 770.1847.

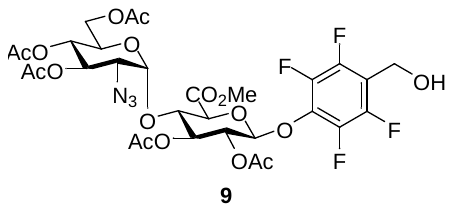

**Methyl (3’,4’,6’-Tri-O-acetyl-2’-azido-2’-deoxy-α-D-glucopyranosyl)-(1→4)-(1-(2,3,5,6-difluoro-1-hydroxy-4-methylhydroxybenzyl)-1- deoxyl-2,3-di-O-acetyl-β-D-glucopyranosyluronate (9):** To a solution of **7** (80 mg, 0.102 mmol) in freshly distilled DCM (2 mL) was added a solution of sodium borohydride (5 mg, 0.122 mmol, 1.2 equiv.) in dry MeOH (200 µL) and the reaction was stirred at room temperature. Conversion of product was monitored by TLC (R*_f_* = 0.32, hexanes/ethyl acetate 1:1). Upon completion, the reaction was quenched by addition of water and extracted into ethyl acetate. The organic layer was washed with brine and dried of Na_2_SO_4_, then purified by FCC using isocratic hexanes/ethyl acetate 1.5/1. White solid product acquired, 51 mg (**64%**).^1^H NMR (600 MHz, Chloroform-*d*) δ 5.37 – 5.29 (m, 2H), 5.24 – 5.19 (m, 3H), 5.02 (dd, *J* = 10.3, 9.3 Hz, 1H), 4.80 (s, 2H), 4.37 (t, *J* = 8.9 Hz, 1H), 4.25 (dd, *J* = 12.5, 3.6 Hz, 1H), 4.13 – 4.07 (m, 2H), 3.85 (ddd, *J* = 10.0, 3.5, 2.0 Hz, 1H), 3.80 (s, 3H), 3.42 (dd, *J* = 10.7, 3.7 Hz, 1H), 2.11 – 2.08 (m, 9H), 2.07 (s, 3H), 2.02 (s, 3H). ^13^C NMR (151 MHz, CDCl_3_) δ 207.17, 170.78, 169.98, 169.76, 169.70, 169.67, 167.28, 146.27, 144.63, 142.02, 141.91, 140.35, 140.24, 134.29, 114.55, 101.64, 98.93, 75.71, 74.48, 73.36, 71.40, 70.24, 68.73, 68.12, 61.25, 61.01, 53.20, 52.82, 36.22, 34.81, 34.67, 31.73, 31.08, 29.84, 29.20, 27.06, 25.42, 22.80, 20.84, 20.82, 20.77, 20.71, 20.68, 18.91, 14.26, 11.57. HRMS (ESI) for C_30_H_33_F_4_N_3_O_17_ [M+Na]^+^ : Calc’d: 806.1643, Found: 806.1662.

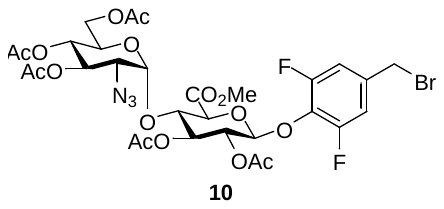

**Methyl (3’,4’,6’-Tri-O-acetyl-2’-azido-2’-deoxy-α-D-glucopyranosyl)-(1→4)-(1-(2,6-difluoro-1-hydroxy-4-methylbromobenzyl)-1- deoxyl-2,3-di-O-acetyl-β-D-glucopyranosyluronate (10):** To a solution of **8** (10 mg, 0.01338 mmol) in freshly distilled DCM (1 mL) was added PPh_3_ (7 mg, 0.0276 mmol, 2 equiv.) and CBr_4_ (9 mg, 0.0276 mmol, 2 equiv.), and the reaction was stirred in the dark at room temperature. Upon complete conversion (monitored by TLC), the solvent was removed *in vacuo* and the crude product was purified by FCC using isocratic hexanes/ethyl acetate 2:1. White solid product acquired, 10 mg (**98%**). ^1^H NMR (600 MHz, Chloroform-*d*) δ 6.97 (d, *J* = 8.1 Hz, 2H), 5.38 – 5.29 (m, 2H), 5.24 – 5.17 (m, 2H), 5.13 (d, *J* = 7.2 Hz, 1H), 5.02 (dd, *J* = 10.3, 9.3 Hz, 1H), 4.37 (s, 2H), 4.35 (d, *J* = 9.1 Hz, 1H), 4.26 (dd, *J* = 12.6, 3.5 Hz, 1H), 4.10 – 4.05 (m, 2H), 3.87 – 3.82 (m, 1H), 3.79 (s, 3H), 3.41 (dd, *J* = 10.7, 3.8 Hz, 1H), 2.09 (d, *J* = 2.0 Hz, 7H), 2.07 (s, 3H), 2.02 (s, 3H). ^13^C NMR (151 MHz, CDCl_3_) δ 170.64, 169.83, 169.61, 169.60, 167.30, 156.16, 154.50, 135.19, 132.44, 113.12, 113.08, 112.96, 101.69, 98.77, 75.71, 74.34, 73.53, 71.52, 70.12, 68.52, 67.98, 61.10, 60.88, 52.98, 31.60, 31.16, 29.71, 22.66, 20.70, 20.64, 20.58, 20.56, 14.13. HRMS spectrum of **X**. HRMS (ESI) for C_30_H_35_F_2_N_3_O_17_ [M+NH_4_]^+^ : Calc’d: 829.1415, Found: 829.1430.

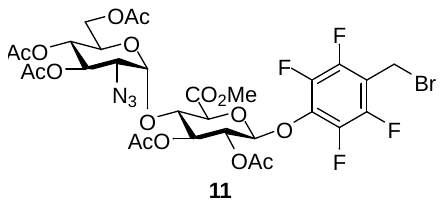

**Methyl (3’,4’,6’-Tri-O-acetyl-2’-azido-2’-deoxy-α-D-glucopyranosyl)-(1→4)-(1-(2,3,5,6-difluoro-1-hydroxy-4-methylbromobenzyl)-1- deoxyl-2,3-di-O-acetyl-β-D-glucopyranosyluronate (11):** To a solution of **9** (10 mg, 0.0128 mmol) in freshly distilled DCM (1 mL) was added PPh_3_ (7 mg, 0.0255 mmol, 2 equiv.) and CBr_4_ (9 mg, 0.0255 mmol, 2 equiv.), and the reaction was stirred in the dark at room temperature. Upon complete conversion (monitored by TLC), the solvent was removed *in vacuo* and the crude product was purified by FCC using isocratic hexanes/ethyl acetate 2:1. White solid product acquired, 10 mg (**99%**). ^1^H NMR (600 MHz, Chloroform-*d*) δ 5.36 – 5.29 (m, 2H), 5.26 (d, *J* = 6.9 Hz, 1H), 5.23 – 5.20 (m, 2H), 5.02 (dd, *J* = 10.3, 9.3 Hz, 1H), 4.49 (d, *J* = 1.3 Hz, 2H), 4.38 (t, *J* = 8.9 Hz, 1H), 4.25 (dd, *J* = 12.6, 3.6 Hz, 1H), 4.15 – 4.11 (m, 1H), 4.09 (dd, *J* = 12.6, 2.2 Hz, 1H), 3.88 – 3.83 (m, 1H), 3.79 (s, 3H), 3.42 (dd, *J* = 10.7, 3.7 Hz, 1H), 2.12 – 2.08 (m, 9H), 2.07 (s, 3H), 2.02 (s, 3H), 1.58 (s, 4H). ^13^C NMR (151 MHz, CDCl_3_) δ 170.79, 169.97, 169.75, 169.74, 167.44, 156.31, 154.64, 135.33, 132.58, 113.26, 113.22, 113.10, 101.83, 98.91, 75.85, 74.48, 73.67, 71.66, 70.26, 68.66, 68.12, 61.24, 61.02, 53.12, 34.82, 31.74, 31.30, 29.85, 29.21, 25.43, 22.84, 22.80, 20.85, 20.78, 20.72, 20.70, 14.27, 11.58. HRMS (ESI) for C_30_H_32_BrF_4_N_3_O_16_ [M+Na]^+^: Calc’d: 869.4783, Found: 870.0763.

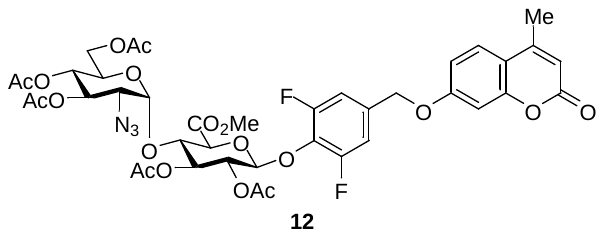

**Methyl (3’,4’,6’-Tri-O-acetyl-2’-azido-2’-deoxy-α-D-glucopyranosyl)-(1→4)-(1-(2,6-difluoro-1-hydroxy-4-methylene(methylumbelliferyl)benzyl)-1- deoxyl-2,3-di-O-acetyl-β-D-glucopyranosyluronate (12):** To a solution of **10** (2 mg, 0.0024 mmol) in dry DMF was added 4-methylumbelliferone (0.6 mg, 0.0036 mmol, 1.5 equiv.) and K_2_CO_3_ (0.8 mg, 0.0048 mmol, 2 equiv.), and the reaction was stirred in the dark at room temperature. Reaction was monitored by HPLC. Upon completion, the product was isolated by HPLC using Method 1. White solid product acquired, 0.5 mg (**25%**). ^1^H NMR (600 MHz, Chloroform-*d*) δ 7.53 (d, *J* = 8.8 Hz, 1H), 7.02 (d, *J* = 8.0 Hz, 2H), 6.92 (dd, *J* = 8.8, 2.5 Hz, 1H), 6.84 (d, *J* = 2.5 Hz, 1H), 6.16 (q, *J* = 1.3 Hz, 1H), 5.38 – 5.30 (m, 2H), 5.25 – 5.19 (m, 2H), 5.14 (d, *J* = 7.2 Hz, 1H), 5.06 (s, 2H), 5.02 (dd, *J* = 10.3, 9.3 Hz, 1H), 4.36 (t, *J* = 9.2 Hz, 1H), 4.26 (dd, *J* = 12.6, 3.5 Hz, 1H), 4.10 – 4.05 (m, 2H), 3.84 (dd, *J* = 10.3, 1.3 Hz, 1H), 3.79 (s, 3H), 3.41 (dd, *J* = 10.7, 3.8 Hz, 1H), 2.41 (d, *J* = 1.3 Hz, 3H), 2.11 – 2.08 (m, 9H), 2.07 (s, 3H), 2.02 (s, 3H). ^13^C NMR (151 MHz, CDCl_3_) δ 171.27, 170.74, 169.93, 169.72, 169.71, 167.41, 161.15, 161.01, 156.65, 156.62, 155.29, 154.99, 154.96, 152.50, 133.91, 133.85, 133.80, 132.42, 132.32, 132.22, 125.93, 114.35, 112.82, 112.56, 111.26, 111.22, 111.13, 111.10, 102.00, 101.92, 98.87, 75.85, 74.46, 73.65, 71.63, 70.21, 68.84, 68.62, 68.08, 61.19, 60.98, 60.51, 53.09, 21.16, 20.81, 20.74, 20.68, 18.79, 14.31. HRMS (ESI) for C_40_H_41_F_2_N_3_O_19_ [M+Na]^+^ : Calc’d: 928.2199, Found: 928.2206.

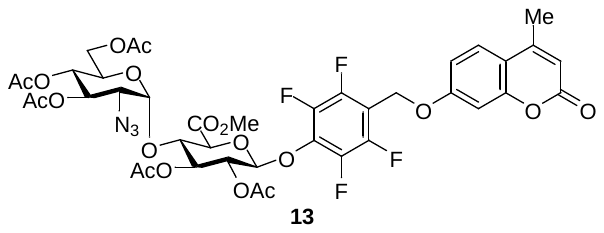

**Methyl (3’,4’,6’-Tri-O-acetyl-2’-azido-2’-deoxy-α-D-glucopyranosyl)-(1→4)-(1-(2,3,5,6-difluoro-1-hydroxy-4- methylene(methylumbelliferyl)benzyl)-1- deoxyl-2,3-di-O-acetyl-β-D-glucopyranosyluronate (13):** To a solution of **11** (5 mg, 0.0059 mmol) in dry DMF was added 4-methylumbelliferone (1.6 mg, 0.0088 mmol, 1.5 equiv.) and K_2_CO_3_ (1.6 mg, 0.0118 mmol, 2 equiv.), and the reaction was stirred in the dark at room temperature. Reaction was monitored by HPLC. Upon completion, the product was isolated by HPLC using Method 1. White solid product acquired, 2.8 mg (**50%**). ^1^H NMR (600 MHz, Chloroform-*d*) δ 7.53 (dd, *J* = 8.7, 4.3 Hz, 1H), 6.96 – 6.86 (m, 2H), 6.18 (q, *J* = 1.3 Hz, 1H), 5.38 – 5.31 (m, 2H), 5.28 (d, *J* = 6.9 Hz, 2H), 5.26 – 5.21 (m, 2H), 5.21 (d, *J* = 4.1 Hz, 2H), 5.18 (s, 2H), 5.03 (dd, *J* = 10.3, 9.3 Hz, 1H), 4.38 (t, *J* = 8.9 Hz, 1H), 4.26 (dd, *J* = 12.6, 3.7 Hz, 1H), 4.17 – 4.06 (m, 3H), 3.85 (ddd, *J* = 10.3, 3.7, 2.2 Hz, 1H), 3.80 (s, 3H), 3.42 (dd, *J* = 10.7, 3.7 Hz, 1H), 2.41 (d, *J* = 1.3 Hz, 3H), 2.11 (s, 6H), 2.10 (s, 9H), 2.09 (s, 6H), 2.07 (s, 4H), 2.03 (s, 3H). ^13^C NMR (151 MHz, CDCl_3_) δ 170.76, 169.98, 169.74, 169.68, 169.64, 167.91, 167.27, 161.18, 160.95, 155.31, 152.47, 146.71, 145.05, 142.05, 140.38, 140.28, 135.31, 132.60, 131.03, 128.95, 125.97, 114.59, 112.74, 112.69, 110.09, 109.97, 109.86, 102.00, 101.51, 98.95, 75.66, 74.49, 73.24, 71.33, 70.24, 68.75, 68.30, 68.12, 61.25, 61.01, 60.54, 58.03, 53.57, 53.22, 38.87, 32.07, 30.50, 29.85, 29.51, 29.07, 23.89, 23.13, 22.84, 21.20, 20.82, 20.81, 20.78, 20.71, 20.69, 18.83, 14.34, 14.26, 14.20, 11.10. HRMS (ESI) for C_40_H_39_F_4_N_3_O_19_ [M+Na]^+^ : Calc’d: 964.2011, Found: 964.1989.

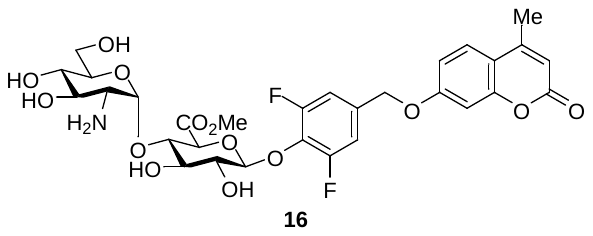

**Methyl (3’,4’,6’-Trihydroxy-2’-amino-2’-deoxy-α-D-glucopyranosyl)-(1→4)-(1-(2,6-difluoro-1-hydroxy-4-methylene(methylumbelliferyl)benzyl)-1- deoxyl-2,3-dihydroxy-β-D-glucopyranosyluronate (16):** To a solution of **11** (0.9 mg, 0.994 x 10^-3^ mmol, 1.0 equiv.) in MeOH (500 µL) was added NaOMe (0.1988 x 10^-3^ mmol, 0.2 equiv.) and the reaction was stirred at room temperature. Reaction progress was monitored by HPLC using Method 1, and upon completion the reaction was quenched by adding DOWEX ion exchange resin until pH ~7. The solvent was removed *in vacuo* and the crude residue was redissolved in MeOH and 10wt% Pd/C added, then charged with hydrogen gas. Reaction stirred at room temperature for 18 h and monitored by HPLC Method 1. Upon completion, the product was purified using HPLC Method 1.

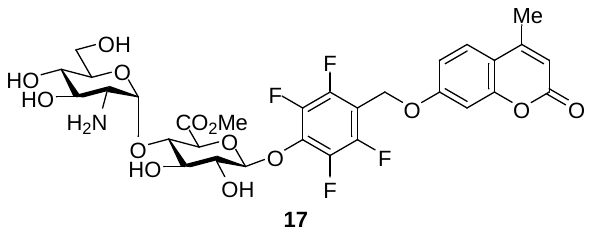

**Methyl (3’,4’,6’-Trihydroxy-2’-amino-2’-deoxy-α-D-glucopyranosyl)-(1→4)-(1-(2,3,5,6-difluoro-1-hydroxy-4- methylene(methylumbelliferyl)benzyl)-1- deoxyl-2,3-dihydroxy-β-D-glucopyranosyluronate (17):** To a solution of **12** (0.9 mg, 0.994 x 10^-3^ mmol, 1.0 equiv.) in MeOH (500 µL) was added NaOMe (0.1988 x 10^-3^ mmol, 0.2 equiv.) and the reaction was stirred at room temperature. Reaction progress was monitored by HPLC using Method 1, and upon completion the reaction was quenched by adding DOWEX ion exchange resin until pH ~7. The solvent was removed *in vacuo* and the crude residue was purified by HPLC using Method 1. The solvent was removed *in vacuo* and the crude residue was redissolved in MeOH and 10wt% Pd/C added, then charged with hydrogen gas. Reaction stirred at room temperature for 18 h and monitored by HPLC Method 1. Upon completion, the product was purified using HPLC Method 1.

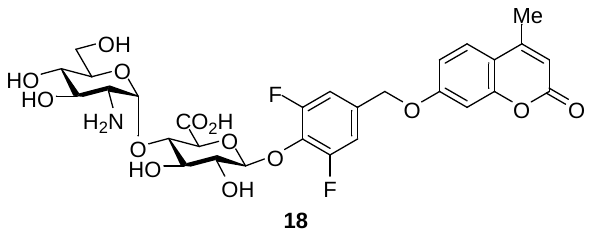

**3’,4’,6’-Trihydroxy-2’-amino-2’-deoxy-α-D-glucopyranosyl)-(1→4)-(1-(2,6-difluoro-1-hydroxy-4-methylene(methylumbelliferyl)benzyl)-1- deoxyl-2,3-dihydroxy-β-D-glucopyranosyluronic acid (18):** SM dissolved in MeOH/H_2_O (1:2) and 1 M NaOH was titrated until reaction pH 12. Stirred at room temperature and monitored by HPLC Method 1. Purified by HPLC Method 1.

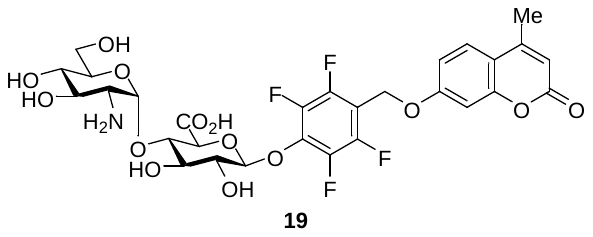

**3’,4’,6’-Trihydroxy-2’-amino-2’-deoxy-α-D-glucopyranosyl)-(1→4)-(1-(2,3,5,6-difluoro-1-hydroxy-4- methylene(methylumbelliferyl)benzyl)-1- deoxyl-2,3-dihydroxy-β-D-glucopyranosyluronic acid (19):** SM dissolved in MeOH/H_2_O (1:2) and 1 M NaOH was titrated until reaction pH 12. Stirred at room temperature and monitored by HPLC Method 1. Purified by HPLC Method 1.

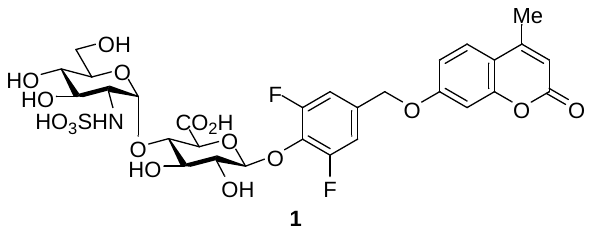

**3’,4’,6’-Trihydroxy-2’-sulfonamido-2’-deoxy-α-D-glucopyranosyl)-(1→4)-(1-(2,6-difluoro-1-hydroxy-4-methylene(methylumbelliferyl)benzyl)-1- deoxyl-2,3-dihydroxy-β-D-glucopyranosyluronic acid (1):** SM dissolved in H_2_O and Py·SO_3_ (10 equiv.) was added. The reaction was stirred at room temperature and monitored by HPLC Method 1.

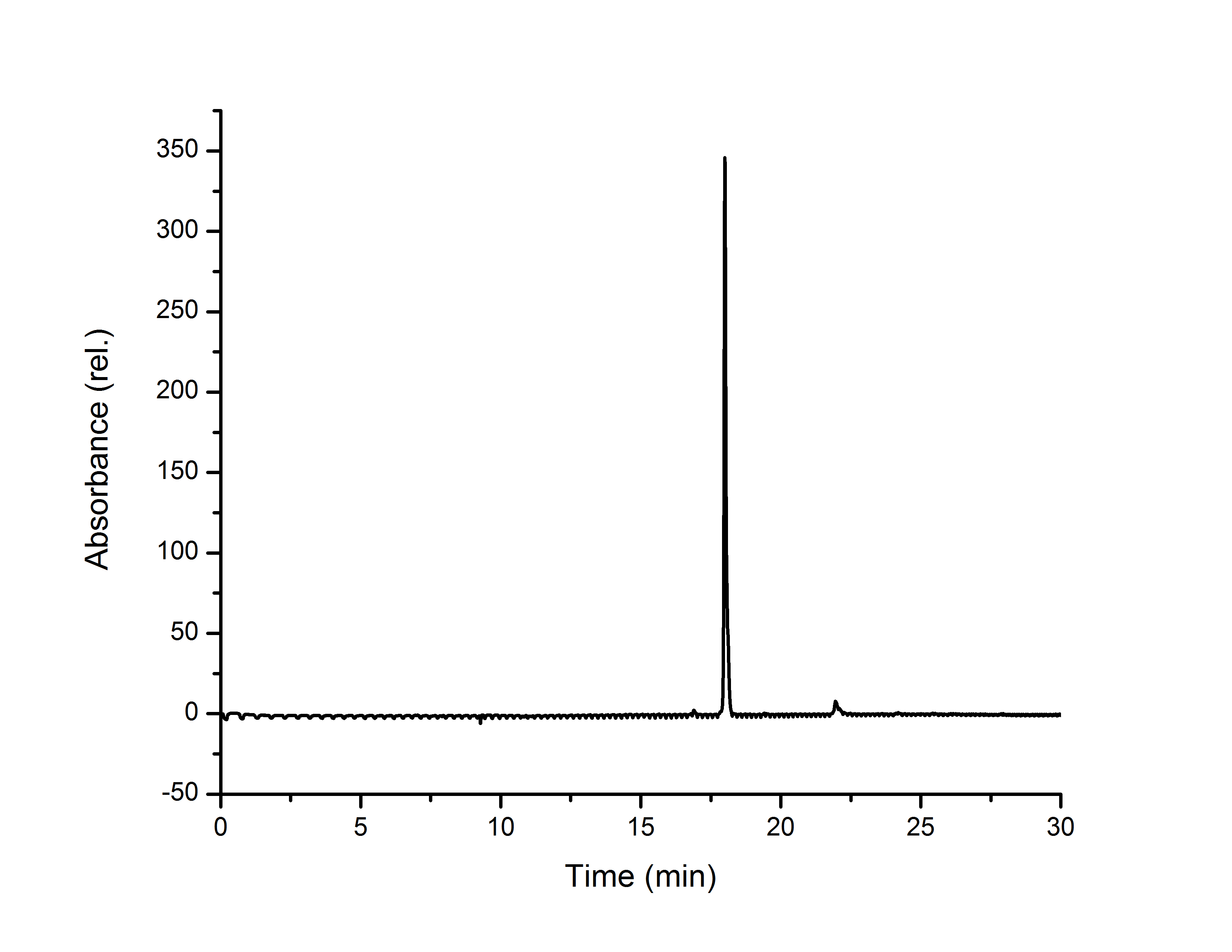

**Figure S1**. HPLC trace of compound **1**, taken using HPLC Method 2.

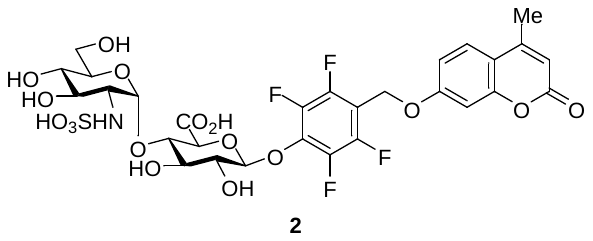

**3’,4’,6’-Trihydroxy-2’- sulfonamido-2’-deoxy-α-D-glucopyranosyl)-(1→4)-(1-(2,3,5,6-difluoro-1-hydroxy-4- methylene(methylumbelliferyl)benzyl)-1- deoxyl-2,3-dihydroxy-β-D-glucopyranosyluronic acid (2):** SM dissolved in H_2_O and Py·SO_3_ (10 equiv.) was added. The reaction was stirred at room temperature and monitored by HPLC Method 1. [M-H]^-^: 770.1 calc’d, 770.3 observed.

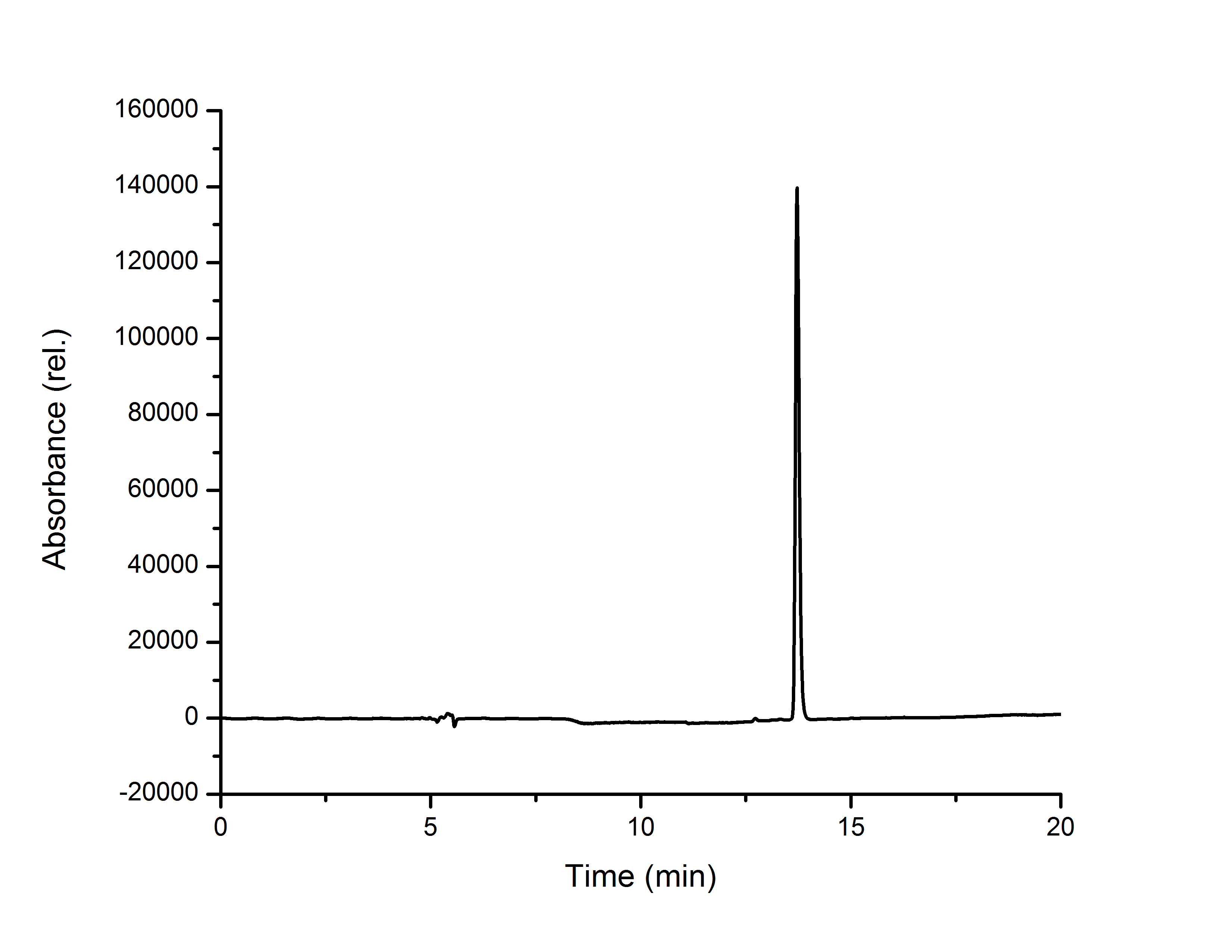

**Figure S2**. HPLC trace of compound **2**, taken using HPLC Method 1.

### Fluorescence assays

#### Fluorescence emission and HPLC assays

Fluorescence assays were run in black-bottom. Probe **2-F** or **4-F** were incubated with HPSE in 40 mM NaOAc buffer, pH 5.0 for 24 h in a 96-well microplate. For **2-F**, the final probe concentration was 35 µM in the presence of 0.4 µg HPSE. For **4-F**, the final probe concentration was 16 µM in the presence of 0.4 µg HPSE, in. All samples were run in duplicate, and fluorescence emission was measured using a BioTek Synergy H1 plate reader. Excitation was set at 365 nm and emission was set at 455 nm. For **4-F** at higher pH, a separate assay was run as described, but after 24 h, 1 M NaOH was added until the assay solution reported a pH of 10. Emission intensity was measured as described. Immediately after analysis, assay wells were injected into HPLC for analysis. For compound **2**, HPLC method 1 was used (Table S1); for compound **1**, HPLC Method 2 (Table S2) was used.

Table S1. HPLC Method 1 – conditions for reaction analysis and purification of compounds.

| Time (min) | Flow (mL/min) | H_2_O (%)^[a]^ | MeCN (%)^[a]^ |
| --- | --- | --- | --- |
| 0 | 3.0 | 98 | 2 |
| 1 | 3.0 | 98 | 2 |
| 11 | 3.0 | 5 | 95 |
| 16 | 3.0 | 5 | 95 |
| 19 | 3.0 | 98 | 2 |
| 21 | 3.0 | 98 | 2 |

Table S2. HPLC Method 2 – conditions for purification of compound 1.

| Time (min) | Flow (mL/min) | H_2_O (%)^[a]^ | MeCN (%)^[a]^ |
| --- | --- | --- | --- |
| 0 | 3.0 | 98 | 2 |
| 1 | 3.0 | 98 | 2 |
| 21 | 3.0 | 5 | 95 |
| 26 | 3.0 | 5 | 95 |
| 29 | 3.0 | 98 | 2 |
| 30 | 3.0 | 98 | 2 |
